# Host-Exclusive Primer Design Enables Effective Metabarcoding via Nanopore Long-Read Sequencing

**DOI:** 10.1101/2024.11.25.625223

**Authors:** Ting He, Koert Jansonius, Xiao Li, Alison M. Reilly, Bahar Sevgin, Rita Setroikromo, Thomas Hackl, Kristina Haslinger

## Abstract

Metabarcoding is a powerful tool to simultaneously identify multiple taxa within a habitat. However, its application to host-associated microbiomes is challenged by substantial co-amplification of host DNA. Here we developed a host-exclusive primer design workflow, to selectively generate amplicons from target taxa while excluding the host. This workflow is centered around a new computational tool, *mbc-prime*, that can generate a list of discriminative candidate primers and score them. We showcase the use of this tool in the design of primers for long-read metabarcoding of endophytic fungi in *Vinca minor. Mbc-prime* streamlines the design of fungus-specific primers, enabling efficient and plant-free amplification of fungal rDNA from mixed DNA samples. Our workflow can be used to study the composition of complex host-associated microbiomes. It should be universally applicable for the design of discriminative primers in a user-friendly and practical manner and thus be of use for various researchers in microbiome research.

## Introduction

High-throughput sequencing of taxonomic marker genes (metabarcoding) has become a widely adopted approach for studying biodiversity and microbial ecology across diverse ecosystems (Abdelfattah *et al*. 2018). Among the critical steps in any metabarcoding workflow, the choice and design of suitable primers is paramount. An optimal primer set must balance broad taxonomic coverage of target organisms while minimizing amplification of non-target taxa (Tedersoo *et al*. 2022).

This issue of non-target amplification extends beyond microbial profiling and is encountered across a wide range of ecological and evolutionary studies. For instance, ancient DNA analyses are often compromised by high levels of microbial or environmental DNA that obscure the target sequences (Slon *et al*. 2017). Marine eDNA studies employing broad-range primers are susceptible to amplification biases, whereby highly abundant eukaryotic taxa, such as dominant zooplankton orders, are preferentially amplified, potentially limiting the detection sensitivity for less abundant taxa within the community (Needham & Fuhrman 2016; Zhan *et al*. 2014).

This complication is particularly pronounced when studying host-associated target taxa, for instance in dietary metabarcoding, where the DNA of the predator or herbivore can dominate over that of the consumed prey or plant material, leading to biased interpretations (De Barba *et al*. 2014); or in parasitological diagnostics where scientists struggle aiming to detect in their hosts frequently struggle with the low abundance of target DNA of helminths or protozoan parasites amidst the overwhelming host background (Avramenko *et al*. 2015). In bacterial profiling of the human microbiome, a common challenge arises from the co-amplification of host organellar DNA that overwhelms the microbial signals, especially in low-biomass niches such as skin, blood, or mucosal surfaces (Karstens *et al*. 2019; Marotz *et al*. 2018). In plant-associated microbiomes, chloroplast and mitochondrial sequences often overshadow bacterial 16S rRNA gene amplification, obscuring true microbial diversity (Beckers *et al*. 2016; Lundberg *et al*. 2013). Fungal profiling, using eukaryotic marker genes such as ITS, 18S, or 28S rRNA, introduces additional complexity. These markers often carry regions that are conserved with host ribosomal DNA, making it difficult to avoid co-amplification, particularly in plant tissues such as roots or leaves (Gdanetz *et al*. 2017; Hadziavdic *et al*. 2014). This can mask the presence of less abundant fungal taxa and complicate downstream ecological interpretation. These examples collectively highlight the widespread impact of non-target DNA interference in amplicon sequencing, emphasizing the need for robust primer design strategies that improve target specificity and minimize the amplification of host DNA to ensure accurate and efficient microbial community profiling.

To overcome these issues, a wide array of bioinformatics tools has been developed to assist in primer design. General-purpose tools like Primer3 (Rozen & Skaletsky 2000), Primer-BLAST (Ye *et al*. 2012), and PrimerQuest™ offer design capabilities based on melting temperature, GC content, and structural properties. However, they are limited in their ability to assess primer suitability for complex, taxonomically diverse samples. Taxon-aware tools such as PrimerProspector (Walters *et al*. 2011) and DegePrime (Hugerth *et al*. 2014) allow for broader coverage assessments and degeneracy control, making them more suitable for environmental applications. Still, these tools generally lack features required for discriminative primer design, such as the ability to target specific taxonomic groups while excluding the host. Other tools, such as BarCrawl (Frank 2009), support barcode design but not primer development; Primer Validator provides taxonomic validation without de novo primer design or advanced scoring systems; and tools like RDP’s Probe Match (Cole *et al*. 2005) or Greengenes’ probe search (DeSantis *et al*. 2006) are limited to database-matching probes without flexible input or customization. Moreover, small-scale tools like Primrose (Ashelford *et al*. 2002) is not scalable to modern large-scale datasets and lack features like 3′-end scoring, which is important for predicting primer success.

Despite this extensive toolkit, most existing programs are not designed to avoid amplification of host DNA, a key requirement for studying host-associated microbiomes. Therefore, there remains a strong need for specialized computational solutions that can design host-exclusive primers, ensuring that sequencing reads derive primarily from microbial taxa rather than host contaminants (Yap *et al*. 2020). Such primers are essential for increasing classification sensitivity, reducing bias, and improving the cost-effectiveness of microbial community profiling in host-rich samples.

Here, we present an effective metabarcoding workflow tailored for host-associated microbiomes, beginning with the design of highly discriminative primers using our new computational tool, *mbc-prime*. This tool automatically identifies and ranks candidate primers based on microbial specificity and host exclusivity. We demonstrate the utility of this approach by profiling the community of endophytic fungi associated with the evergreen plant *Vinca minor* (lesser periwinkle), using long-read nanopore sequencing. Our method provides a robust solution for the accurate detection of microbial and other organismal diversity in challenging, host-associated environments.

## Materials and Methods

### Sample collection and DNA extraction from the leaves of *Vinca minor*

For the sampling, 10 - 15 individual plants of *Vinca minor* were collected from Italy (45°26’3’’ N, 8°49’53’’W), immediately placed in an ice box, and transported to the lab. After surface sterilization (He *et al*. 2023), the plant material was quickly frozen with liquid nitrogen, ground into fine powder, and stored at – 70 ℃ until DNA extraction.

Holobiont genomic DNA was extracted using NucleoSpin® Microbial DNA kit (BIOKÉ, Leiden, The Netherlands) in accordance with the protocols provided by the manufacturer with adjustment. Briefly, 0.2 g ground plant material was suspended in 100 µL elution buffer BE, 40 µL buffer MG and 10 µL liquid proteinase K. The plant cells were disrupted by vortexing at 1000 rpm for 3 mins. After adding 600 µL buffer MG, samples were vortexed for another 1 min at 1000 rpm, followed by 2 min centrifugation at 13,000 x g. The supernatant was transferred onto a NucleoSpin® Microbial column, washed twice with buffer BW and B5 buffer, and eluted with 30 µL elution buffer BE. The DNA was quantified with NanoDrop N-100 (ThermoFisher, Waltham, MA, USA) and stored in – 20 ℃ until further use.

### Mock community construction

To generate a synthetic mock community for initial testing of the new primers, we used six species isolated from healthy-looking, surface-sterilized leaves of *Vinca minor,* collected in Groningen (The Netherlands) in November 2021. Fungal isolation, cultivation, and genomic DNA isolation were executed as described in our previous work (He *et al*. 2023). The SR1R-Fw/ LR12-R primer pair (D’Andreano *et al*. 2021) was used to amplify the entire fungal rDNA locus. Reactions were run in volumes of 25 µL, containing 1 μL each of the forward (10 µM) and reverse primer (10 µM), 12.5 µL LongAmp Hot Start Taq 2x Master Mix (New England Biolab, MA, USA), 1 µL of DNA template (40 fmol), and 9.5 µL of nuclease-free water. PCR amplification was carried out with the following protocol: initial denaturation of 94°C for 30 s, followed by 30 cycles of 94°C for 10 s, 65°C for 30 s, and 65°C for 5.5 min, followed by a final 65°C extension for 10 min. The PCR products were purified using AMPure XP magnetic beads (Beckman Coulter, California, USA) and the concentrations were determined using Qubit 3.0 (Invitrogen, Waltham, MA, USA), and the Qubit dsDNA HS Assay Kit. These six fungal amplicons were mixed in equimolar ratios to construct the mock community DNA sample with a concentration of 55 ng/ µL.

### Primer design and implementation on endophytic fungi in *Vinca minor*

To analyze endophytic fungi in *Vinca minor*, all related sequences were downloaded from the SILVA database (Supplementary File 2, Datasets S1 and S2). For the SSU region, 46 sequences from the Gentianales order were extracted from the downloaded database file (SILVA_138.1_SSURef_NR99_tax_silva.fasta.gz.). In the same way, a dataset of 118 fungal sequences was collected, which represents all major fungal divisions (ascomycota, basidiomycota, chytridiomycota, cryptomycota, zoopagomycota, microsporidia, mucoromycota, neocallimastigomycota and blastocladiomycota). These 164 sequences were aligned using the MAFFT multiple sequence alignment software (ouput format: fasta format, input order; strategy: L-INS-i; arguments: --ep)(Katoh & Standley 2013). For the LSU region, 62 fungal sequences (Supplementary File 2, Dataset S3) and 6 available Gentianales sequences (Supplementary File 2, Dataset S4) were extracted from the downloaded database file (SILVA_138.1_LSURef_NR99_tax_silva.fasta.gz). Alignment was performed among these 68 sequences in the same way as for the SSU.

Both alignments were analyzed with *mbc-prime* version 0.6.0 (-s = 0.5) (see Results section for implementation). We obtained 147 candidate discriminative primers across the SSU (Supplementary File 2, Dataset S5) and 371 across the LSU (Supplementary File 2, Dataset S6). Four candidates for forward primers on the SSU and two reverse primers on the LSU were selected from the initial set based on the following criteria: 1) at least 80% coverage of fungi at zero or one mismatch (*in silico*); 2) at least one mismatch at the 3’ end between primer compared to the plant sequences; 3) a GC content between 40 and 60%; 4) approx. 20 base pairs in length. In addition, we manually optimized the length of candidate primer sequences to lower the difference in melting temperatures and the likelihood of heterodimer formation in primer combinations as well as incorporated degenerate positions to maximize the coverage of the target group. Finally, all selected primers were evaluated *in silico* using SILVA TestProbe 3.0 (https://www.arb-silva.de/search/testprobe/) for the individual primers and OligoAnalyzer™ (Version 3.1, Integrated DNA Technologies, https://eu.idtdna.com/pages/tools/oligoanalyzer) for primer pairs before being tested on the DNA samples of a fungal mock community and plant material. A step-by-step tutorial for using *mbc-prime* is listed in the Supplementary Note S1.

### Amplicon generation for Oxford Nanopore sequencing

The strategy employed in this study involves two steps of PCR. The first PCR was performed with the gene-specific rDNA primers including overhangs that serve as handles for PCR barcoding in the second step (5’ TTTCTGTTGGTGCTGATATTGC-[forward primer sequence] 3’; 5’ ACTTGCCTGTCGCTCTATCTTC-[reverse primer sequence] 3’). All primer pairs used in this study are shown in Table 2 and sequences of all primers used in this study are listed in Supplementary File 2, Dataset S7. Each PCR reaction mix consisted of 0.4 μM each of the forward and reverse primer, 1 x LongAmp Hot Start Taq 2x Master Mix (New England Biolab, MA, USA), and 40 ng of template DNA, combined in a 25 µL reaction. The PCR cycling conditions were as follows: initial denaturation of 94°C for 30 s, followed by 30 cycles of a 10 s hold at 94°C, a 30 s hold at the respective annealing temperature shown in Table 1, and an elongation period specific for the length of the amplicon as shown in table 1 at 65°C, followed by a final 65°C extension for 10 min. The PCR amplicons were purified using AMPure XP magnetic beads (Beckman Coulter, California, USA) and quantified by Qubit. The second PCR was used to incorporate the Oxford Nanopore barcode sequences with the PCR Barcoding Expansion 1-12 (EXP-PBC001, ONT, Oxford, UK). Each amplicon PCR product was used as a template (100-200 fmol) in the second PCR reaction with 1 µL PCR barcoding primer mix (one of BC1-BC12, at 10 µM), and 25 µL LongAmp Hot Start Taq 2X Master Mix (New England Biolab, MA, USA) in a total of 50 µL. The PCR cycling conditions were as follows: initial denaturation of 95°C for 3 min, followed by 15 cycles of 95°C for 15 s, 62°C for 15 s, and an elongation period specific for the length of the amplicon as shown in table 1 at 65°C, followed by a final 65°C extension for 10 min. The barcoded PCR products were further purified with 0.4x volume of AMPure XP Beads (Beckman Coulter, California, USA). All barcoded PCR products were pooled in equal amounts to prepare 1 µg of DNA in 50 µL nuclease-free water for library preparation. The metadata of samples as input for the library preparation is listed in the Supplementary File 2, Dataset S8.

**Table 1:**
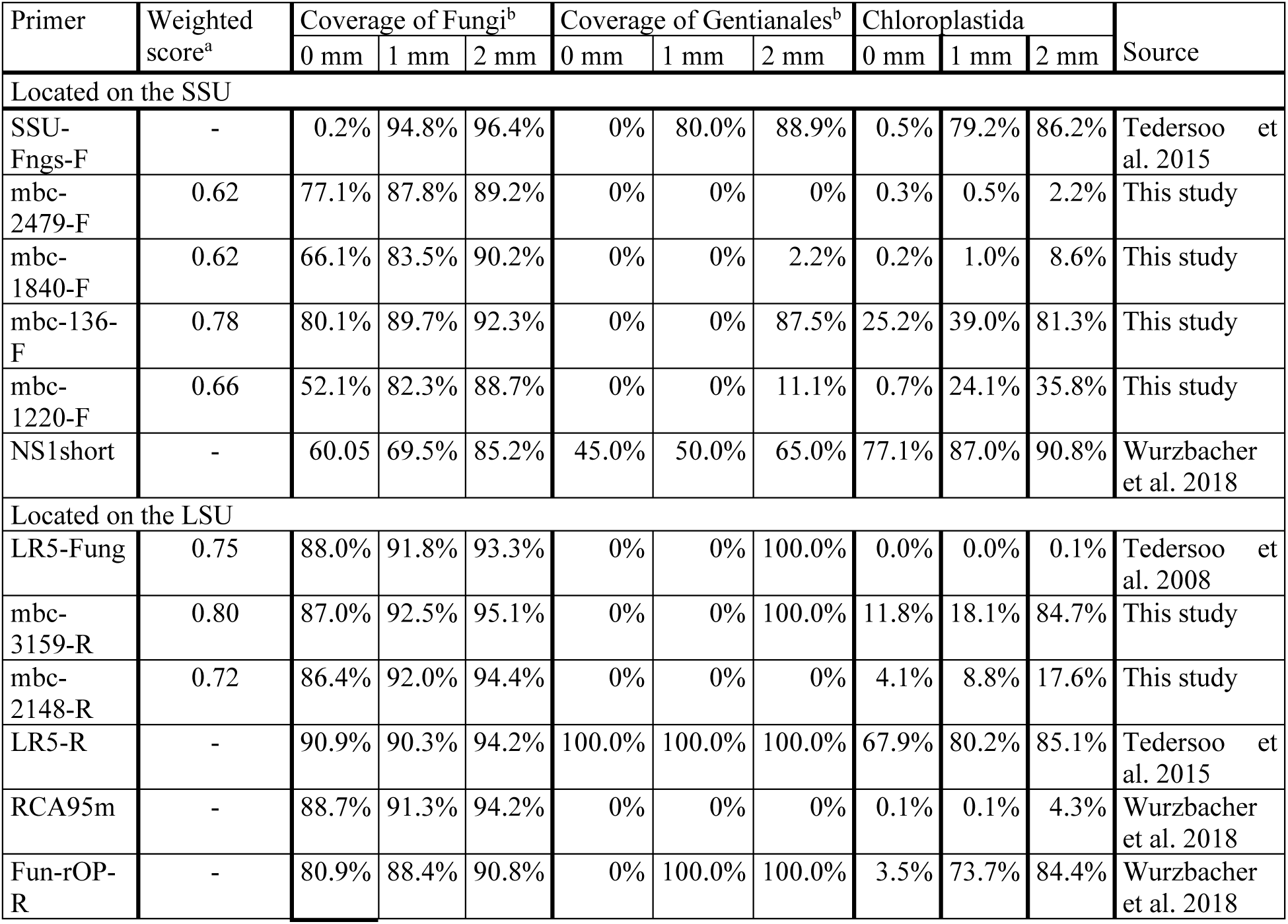
*mbc-prime* and SILVA TestProbe 3.0 evaluations of all primers used in this study. a: *mbc-prime* weighted score of the candidate primer before manual modification (optional); b: summative coverage achieved with the final primers determined by SILVA TestProbe 3.0; mm: number of maximum mismatches

| Primer | Weighted score<sup>a</sup> | Coverage of Fungi<sup>b</sup> |  |  | Coverage of Gentianales<sup>b</sup> |  |  | Chloroplastida |  |  | Source |
| --- | --- | --- | --- | --- | --- | --- | --- | --- | --- | --- | --- |
|  |  | 0 mm | 1 mm | 2 mm | 0 mm | 1 mm | 2 mm | 0 mm | 1 mm | 2 mm |  |
| Located on the SSU |  |  |  |  |  |  |  |  |  |  |  |
| SSU-Fngs-F | - | 0.2% | 94.8% | 96.4% | 0% | 80.0% | 88.9% | 0.5% | 79.2% | 86.2% | Tedersoo et al. 2015 |
| mbc-2479-F | 0.62 | 77.1% | 87.8% | 89.2% | 0% | 0% | 0% | 0.3% | 0.5% | 2.2% | This study |
| mbc-1840-F | 0.62 | 66.1% | 83.5% | 90.2% | 0% | 0% | 2.2% | 0.2% | 1.0% | 8.6% | This study |
| mbc-136-F | 0.78 | 80.1% | 89.7% | 92.3% | 0% | 0% | 87.5% | 25.2% | 39.0% | 81.3% | This study |
| mbc-1220-F | 0.66 | 52.1% | 82.3% | 88.7% | 0% | 0% | 11.1% | 0.7% | 24.1% | 35.8% | This study |
| NS1short | - | 60.05 | 69.5% | 85.2% | 45.0% | 50.0% | 65.0% | 77.1% | 87.0% | 90.8% | Wurzbacher et al. 2018 |
| Located on the LSU |  |  |  |  |  |  |  |  |  |  |  |
| LR5-Fung | 0.75 | 88.0% | 91.8% | 93.3% | 0% | 0% | 100.0% | 0.0% | 0.0% | 0.1% | Tedersoo et al. 2008 |
| mbc-3159-R | 0.80 | 87.0% | 92.5% | 95.1% | 0% | 0% | 100.0% | 11.8% | 18.1% | 84.7% | This study |
| mbc-2148-R | 0.72 | 86.4% | 92.0% | 94.4% | 0% | 0% | 0% | 4.1% | 8.8% | 17.6% | This study |
| LR5-R | - | 90.9% | 90.3% | 94.2% | 100.0% | 100.0% | 100.0% | 67.9% | 80.2% | 85.1% | Tedersoo et al. 2015 |
| RCA95m | - | 88.7% | 91.3% | 94.2% | 0% | 0% | 0% | 0.1% | 0.1% | 4.3% | Wurzbacher et al. 2018 |
| Fun-rOP-R | - | 80.9% | 88.4% | 90.8% | 0% | 100.0% | 100.0% | 3.5% | 73.7% | 84.4% | Wurzbacher et al. 2018 |

### Library preparation

The library preparation was performed with the ligation sequencing kit V14 (ONT, SQK-LSK114) following the manufacturer’s instructions (https://community.nanoporetech.com/docs/prepare/library_prep_protocols/ligation-sequencing-gdna-pcr-barcoding-sqk-lsk114-with-exp/v/pbc_9182_v114_revh_07mar2023). First, the pooled barcoded amplicons underwent end repair with the NEBNext® Ultra II End Repair / dA-tailing Module (NEB, E7546). This reaction consisted of 50 µL DNA (1 µg), 7 µL Ultra II End-prep Reaction Buffer and 3 µL Ultra II End-prep Enzyme Mix, and was incubated at 20°C for 5 minutes and 65°C for 5 minutes. The end-repaired amplicons were further purified using AMPure XP magnetic beads and quantified by Qubit. For the adapter ligation step, 60 µL of the end-repaired and purified amplicons, 25 µL Ligation Buffer (LNB), 10 µL NEBNext Quick T4 DNA Ligase, and 5 µL Ligation Adapter (LA) were combined in a reaction tube and incubated for 10 min at room temperature. After ligation, the samples were mixed with 0.4 x volume of AMPure XP magnetic beads and then washed twice with 250 µL Long Fragment Buffer provided in the ligation sequencing kit. The library was finally eluted with 15 µL Elution Buffer (EB), incubated at 37 °C for 10 minutes, and quantified by Qubit. 35-50 fmol of this final prepared library were loaded for sequencing.MinION sequencing data analysis Sequencing was performed on a MinION device with a R10.4.1 flow cell running with MinKNOW software v24.02.10 . Basecalling and demultiplexing were performed with Dorado v7.3.9 in super accurate mode. We trimmed reads using NanoFilt v2.8.0 and assessed the read quality using NanoPlot v1.42.0. Subsequently, the FASTQ files were uploaded to EPI2ME Labs v4.1.4, and the “wf-metagenomics” workflow was performed with the following settings: classifier: Kraken2; Kraken2 memory mapping: true; dataset: ncbi_16s_18s_28s_ITS or SILVA_138_1; N taxa barplot: 9; bracken length: 2000; min percent identity: 90; min len: 2000; max len: 6000. The mock community samples were analyzed with a custom database built with Kraken2. Relative abundances were further visualized with GraphPad Prism 10.

## Results

### Rationale and workflow for effective host-exclusive primer design

Metabarcoding relies on the targeted PCR amplification of suitable genomic markers that comprise conserved regions serving as the primer binding sites and variable regions with enough differences to distinguish individual species. Herein, it is crucial that the primers amplify DNA equally well across a broad taxonomic range. Different marker genes are used for different biological groups: for animals, the mitochondrial gene cytochrome c oxidase I (COI) is widely applied (Hebert *et al*. 2003); in plants, chloroplast genes such as *matK* and *rbcL* are commonly used (Hollingsworth *et al*. 2009); for bacteria and archaea, the 16S rRNA gene is the standard marker, while fungi and protists are typically identified using the internal transcribed spacer (ITS) regions, along with the 18S and 28S rRNA genes (Klindworth *et al*. 2013).

Expanding on this principle, the rationale for creating host-exclusive, discriminative primers is simple: identify a variable-region-enclosing primer pair that is conserved among all target group members but differs in sequence in all unwanted taxa. Such locations can be discerned in one or more combined multiple-sequence alignments (MSAs) of representative sequences of the target and exclusion group. This rationale allowed us to devise an effective workflow for host-exclusive primer design for holobiont metabarcoding (Figure 1). Since this workflow requires identifying discriminative primer sites in MSAs, which is slow and tedious if done by hand, we also devised the program *mbc-prime* (described in detail in the next section) to help automate this crucial step. The list of discriminative, candidate primers returned by *mbc-prime* can then be further inspected for desired properties of the primer, for instance the GC content and melting temperature of the primer and the position of the non-target mismatches within the primer sequence. With the most discriminative primers in hand, additional optimization of the primers can be performed by introducing degenerate bases into the sequence to optimize the coverage among the target sequences. Lastly, the candidate primers can be evaluated for their specificity for target group and efficiency of excluding non-target group.

**Figure 1.**
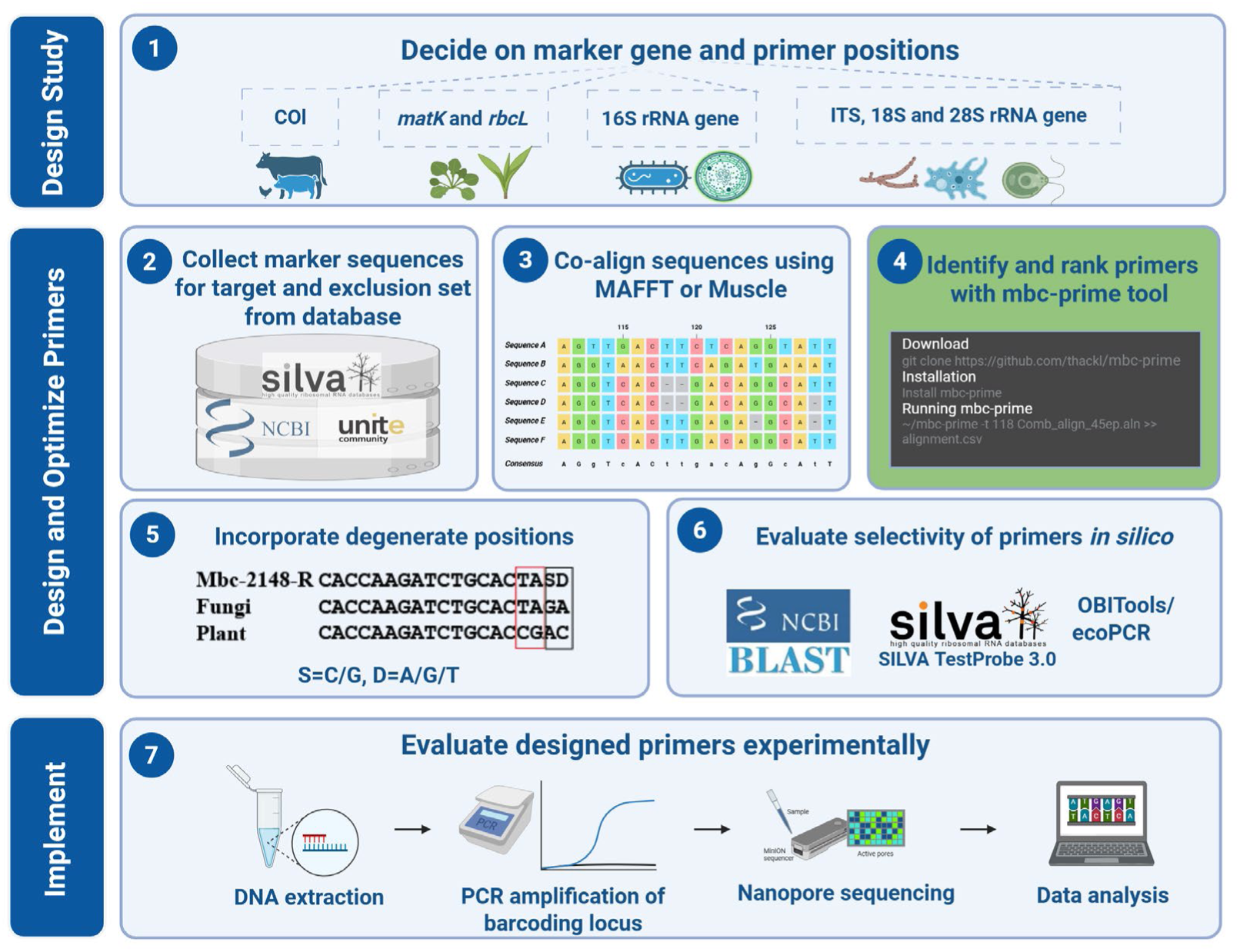
Workflow for effective host-exclusive primer design as described in this study. At the core of the workflow stands the computational tool mbc-prime, which effectively identifies candidates for discriminative primers based on multiple sequence alignments of the target and the exclusion sets (step 4).

### Detection of discriminative primer binding sites with *mbc-prime*

Discriminative primers should align optimally with all target sequences while differing maximally from all non-target sequences. To identify such primers computationally, we developed *mbc-prime*, a stand-alone Python program available at https://github.com/thackl/mbc-prime.

*mbc-prime* identifies discriminative primers based on a user-supplied MSA in FASTA format, comprising DNA sequences of the chosen genomic locus in the target and exclusion sets. In a first step, the MSA is imported and preprocessed. Terminal gaps, introduced by partial sequences frequently present in amplicon marker databases, are masked to distinguish them from genuine gaps. Columns with more than 20% (default value) gaps or ambiguous bases (non-ATGCU) are trimmed from the alignment. The trimmed MSA is then analyzed via a sliding window with a size corresponding to the intended primer length (default: 20 bp). For each window and sequence set (target and exclusion), the abundance of each sequence variant is counted, and the most abundant sequence variant of the target set is chosen as primer candidate. Next, for each remaining sequence variant from both sets, the number of mismatches compared to the primer candidate is computed, and the overall percentage of sequence variants with 0, 1, 2, 3 or 4-or-more mismatches are determined. The underlying edit distances are computed via Python bindings of the Edlib library (Šošić & Šikić 2017). This results in two mismatch profiles per primer candidate, one for the target set and one for the exclusion set. An example of promising profiles for a discriminative primer would be the following: 90% and 10% of the target sequences align with 0 and 1 mismatch, respectively, while 10%, 20% and 70% of the exclusion sequences align with 2, 3, and 4-or-more mismatches, respectively.

To facilitate easy comparison and ranking of possible primer candidates based on their mismatch profiles, we integrate the two profiles obtained from target and exclusions sets into a single discriminative score, as defined below. The score ranges between 0 and 1, with 1 capturing 100% of target sequences perfectly while excluding 100% of non-target sequences at the given mismatch threshold (default mismatch threshold: mm = 2).

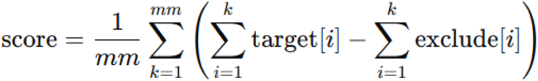

Finally, candidate primers in the MSA are filtered for a maximum number of gaps, ambiguous bases and a minimum score (default 0.5), and overlapping primers are grouped together. The filtered list of above-threshold primer candidates is reported together with the positional information, score, mismatch profiles, and sequence in a tab-separated table (Figure 2).

**Figure 2.**
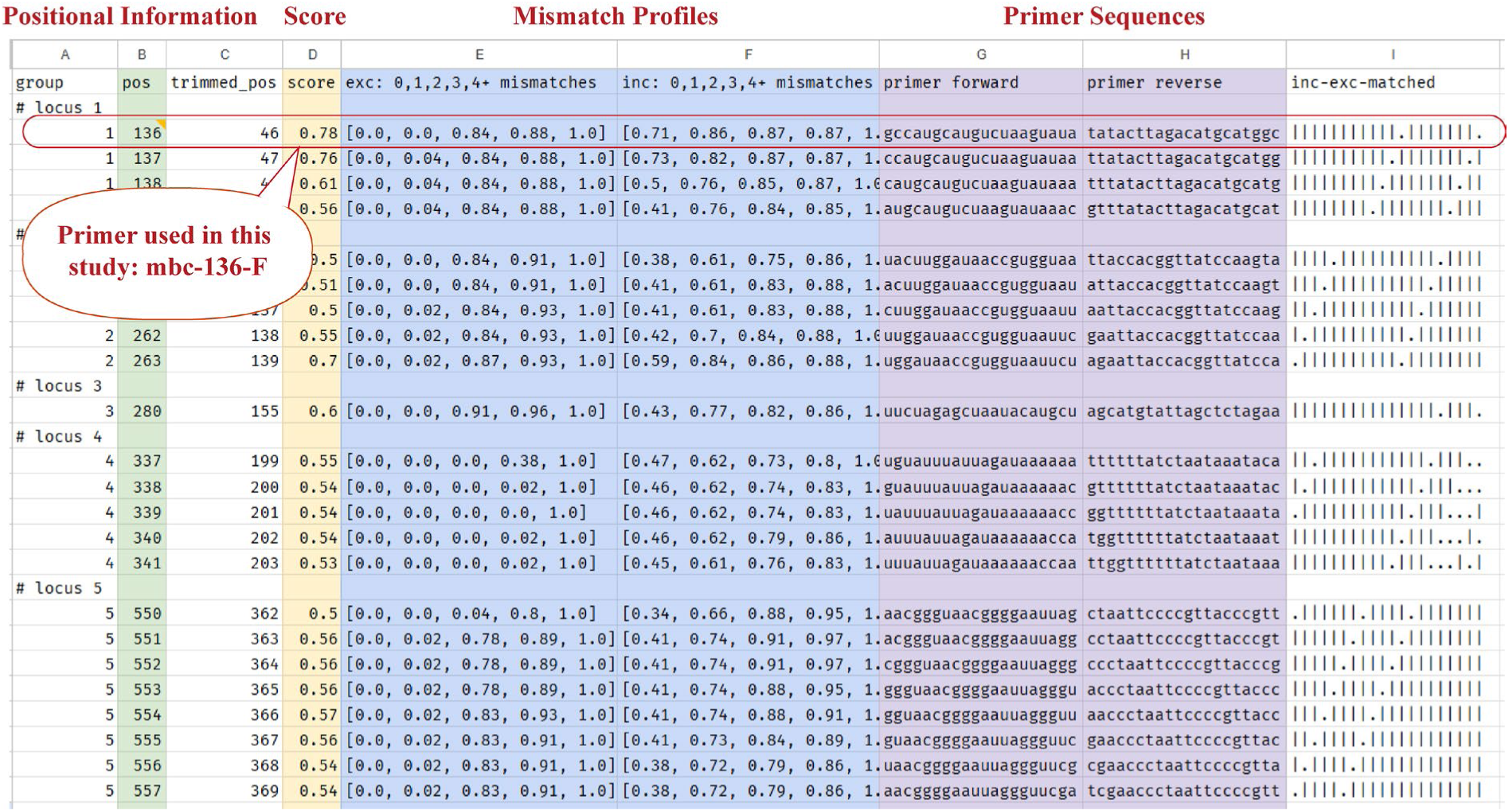
Screen shot of an example output file of mbc-prime.

### Case study

#### Design of metabarcoding primers for endophytic fungi in *Vinca minor*

To demonstrate the applicability of our metabarcoding primer design workflow, we targeted the endophytic mycobiome of *Vinca minor* (lesser periwinkle) with long-read sequencing of the rDNA locus.

*V. minor* belongs to the order of Gentianales in the Apocynaceae family and is a popular plant for its ornamental and medicinal properties. To date, the microbiome of *V. minor* remains largely unexplored. In our initial amplicon sequencing experiments with published primers targeting the rDNA of fungi for long-read sequencing, we observed that 99 % of reads were derived from the *V. minor* rDNA locus (data not shown). Therefore, we considered this the ideal test case for the performance of our holobiont metabarcoding workflow.

Since we wanted to target the entire rDNA locus, we downloaded fungal (target) and plant (exclusion) sequences spanning the SSU and LSU regions from the SILVA database. We selected 118 fungal SSU sequences from different phyla (Ascomycota, Basidiomycota, Chytridiomycota, Cryptomycota, Zoopagomycota, Microsporidia, Mucoromycota, Neocallimastigomycota, and Blastocladiomycota) to represent all major fungal divisions (Supplementary File 2, Dataset S1), and 46 sequences from the order Gentianales (Supplementary File 2, Dataset S2). For the LSU, we included 62 fungal sequences (Supplementary File 2, Dataset S3) and 6 sequences from Gentianales (Supplementary File 2, Dataset S4). We created two MSAs for all SSU and LSU sequences, respectively, and scanned them for discriminative primer binding sites. Using *mbc-prime* with default settings, we obtained 147 candidate primers across the SSU (Supplementary File 2, Dataset S5) and 371 across the LSU (Supplementary File 2, Dataset S6). From the list, we selected primers that had scores of 0.6 and higher and carried at least one mismatch at the 3’ end when compared to plant (exclusion) sequences. With these criteria we obtained four forward primers in different regions of the SSU and two reverse primers in different regions of the LSU (Figure 3A). Following careful examination of the GC content, the predicted melting temperatures, and the respective primer binding sites within the MSA, we manually optimized the primer sequences to establish the final sequences (Figure 3B, Supplementary File 2, Dataset S7). Primer mbc-1220-F was extended by two nucleobases at the ‘5 end, mbc-2148-R was shortened by one base at the ‘5 end, and in mbc-2479-F and mbc-2148-R we incorporated degenerate positions to maximize the coverage of the target group (Fig. S1).

**Figure 3.**
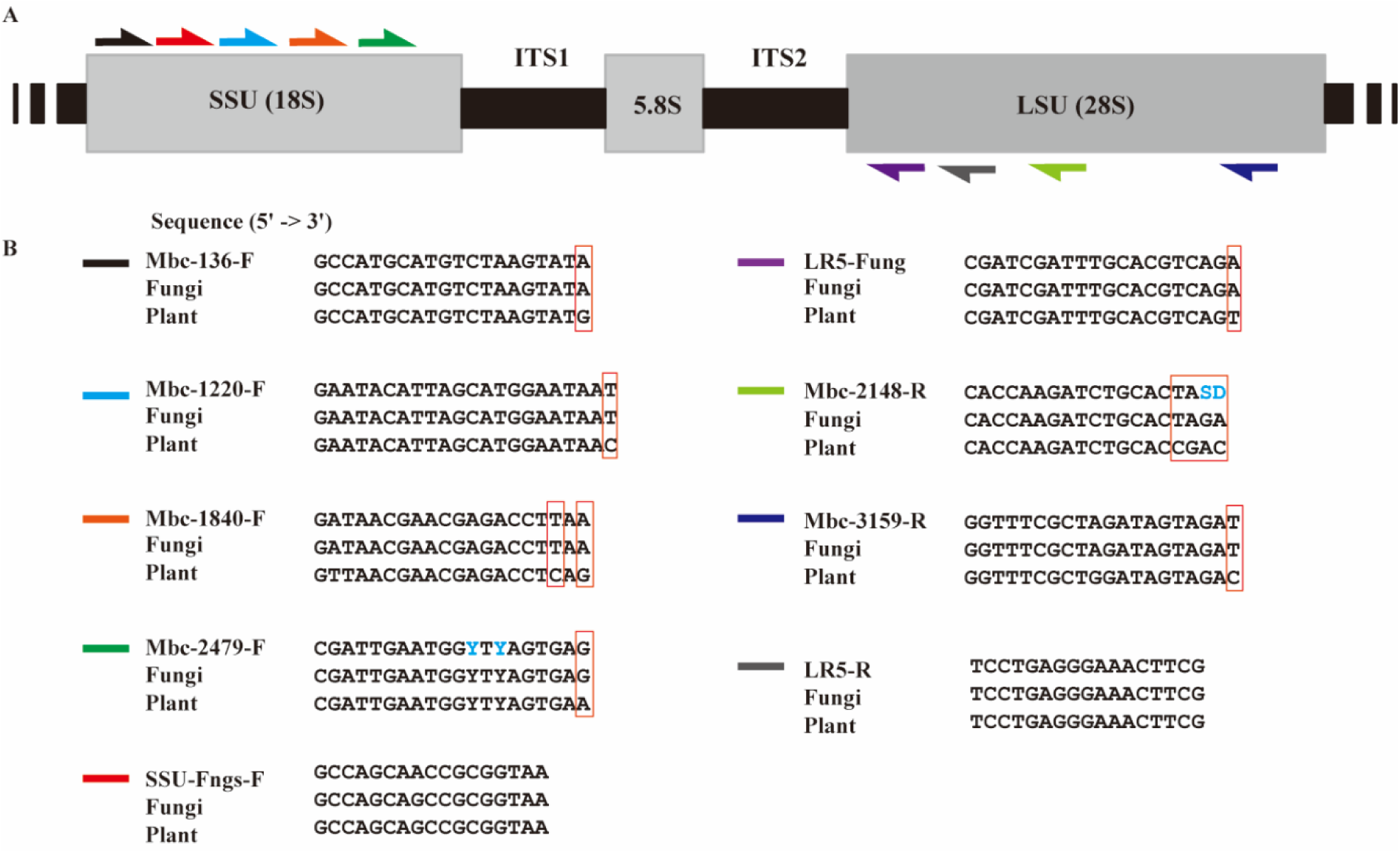
A. Illustration of primer locations in the rDNA locus consisting of SSU, ITS1, 5.8S, ITS2, and LSU; B. Sequence alignments of primers with the consensus sequences of fungi (target set) and plant (exclusion set). Degenerate bases (marked in blue): Y = C/T, S = C/G, D = A/G/T.

To assess the performance of the final primers *in silico*, we used the SILVA TestProbe 3.0 tool with default settings and “fungi” and “Gentianales” or “Chloroplastida” as test sets. For completeness, we also included several primers from literature (Nilsson *et al*. 2019; Tedersoo *et al*. 2008, 2015; Wurzbacher *et al*. 2019) Similar to the literature primers, all of the new primers show a wide coverage across fungi. In terms of coverage across the Chloroplastida, our newly designed primers, similar to the literature primers LR5-Fung and RCA95m, exhibited low coverage within this group (Table 1). We included LR5-Fung, which is also identified by our *mbc-prim*e tool, for further experimental analysis.

With these primer candidates in hand, we performed an *in silico* analysis of all possible primer combinations to identify those with favorable parameters, such as a low difference in melting temperatures and a low likelihood of heterodimer formation. The 10 primer pairs that met our quality standards (pairs 1-10; Table S1, Table S2) and one primer pair from literature (“3.5 kb”) (D’Andreano *et al*. 2021) were carried forward for experimental validation with a fungal mock community and genomic DNA extracted from the surface-sterilized *V. minor* holobiont.

#### Experimental evaluation of the designed fungus-specific primers using mock community

We generated a standardized DNA sample by mixing equimolar amounts of PCR-amplified rDNA from six fungal isolates of our in-house strain collection (Table S3). The strains were chosen to cover a wide taxonomic range and different levels of GC content in order to assemble an authentic mock community for our primer testing. Next, we optimized the annealing temperatures of the PCR for each primer pair to generate high-quality amplicons based on visual inspection of the bands after agarose gelelectrophoresis (Figure 3A). All primer pairs yielded clear bands of the appropriate molecular weight. Since it was our primary goal to generate long amplicons covering both SSU and LSU, we abandoned primer pairs 1-6 after this step and proceeded with multiplex sequencing of the amplicons generated with primer pairs 7-10 with the R10.4.1 flow cells from Oxford Nanopore Technologies. All steps from PCR to sequencing were performed with technical duplicates and in parallel to a positive control (3.5 kB primer pair) and a negative control (nuclease-free water). With one sequencing run, we generated 303,100 raw reads with a yield of 1.1 Gb and a mean Qscore of 18 (accuracy: 98%; Table S4).

To classify the trimmed and filtered reads, we applied the *wf-metagenomics* workflow with Kraken2. Since not all target regions of the mock community members were available in public databases, we generated a custom database comprising the whole genome sequences of the six isolates with Kraken2 (Supplementary Note S2). Using this custom database, 99.80% of reads across all samples were classified, indicating a high sample quality with minimal contamination or sample preparation artefacts, and a high sequencing accuracy. With regards to the relative abundances of species within the mock community, all samples showed a similar composition although among the amplicons generated with primer pair 8, the basidiomycete *Trametes versicolor* seemed to be underrepresented. This indicates that all primer pairs except for primer pair 8, successfully amplify rDNA regions across diverse fungal isolates, producing amplicons that closely match the expected composition of the mock community (Figure 4B).

**Figure 4.**
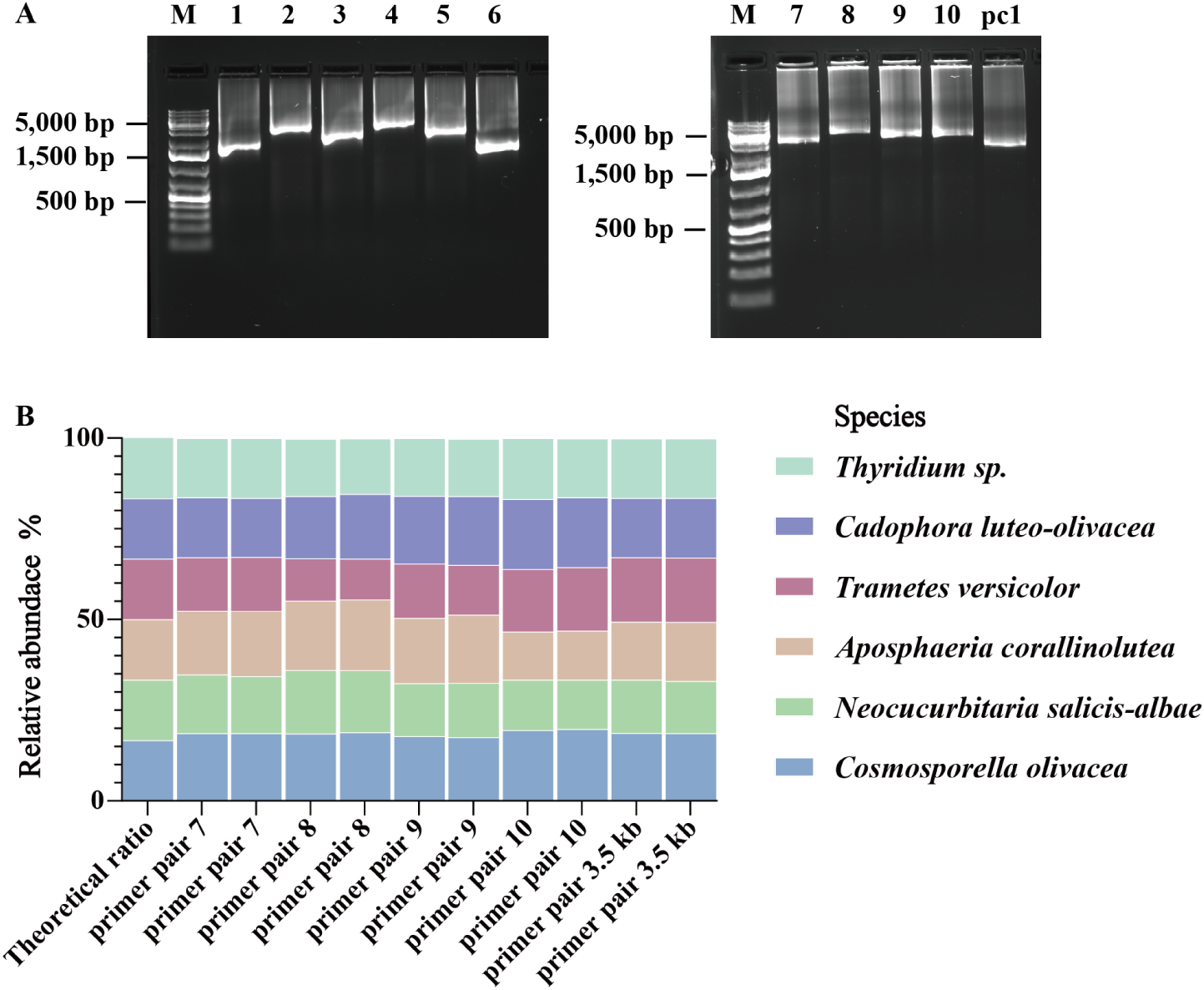
Metabarcoding of a fungal mock community. (A) Gel electrophoresis of rDNA amplicons generated from gene specific primers including overhangs for PCR barcoding; Sample order: M: 1 kb plus DNA ladder; 1-10: amplicons generated with primer pairs 1-10 respectively; pc1: positive control generated with primer pair 3.5 kb. All primer pairs yielded products in the expected size range. (B) Relative abundance of taxonomic groups classified based on the sequencing reads of amplicons generated from the mock community with the different primer pairs (each primer pair has two replicates). Species belonging to the same family are displayed in the same color with different fill patterns. All primer pairs reproduce the overall structure of the mock community, albeit with different biases for different groups.

#### Experimental evaluation of the designed fungus-specific primers using genomic holobiont DNA of *V. minor*

Having confirmed the amplification efficacy of the new primer pairs on the mock community sample, we next sought to assess their performance in a real-world holobiont scenario. We used the primer pairs to specifically target the endophytic fungi in the holobiont DNA from *V. minor*. However, while preparing for the experiment and performing the first amplification step with the holobiont DNA (Figure S3), we noticed that primer pair 8 did not give a high-quality PCR product with a distinct band of the correct size. The individual primers performed well in the other primer combinations thus we conclude that the issue specifically arises from the combination of these primers in the context of the holobiont sample, where the target fungal DNA is rare. Since we could not overcome this problem despite various optimization attempts (annealing temperature, additives, polymerase), we abandoned primer pair 8.

Proceeding with the remaining three discriminative primer pairs 7, 9, and 10, and the non-discriminative literature-based primer pair 3.5kb, we generated, a total of 599,335.274 reads with a Qscore greater than 18 in a single multiplex sequencing run (Table S5). When using the same classification approach as before (*wf-metagenomics* workflow, Kraken2), this time with a custom database comprising the NCBI RefSeq Targeted Loci database (ncbi_16s_18s_28s_ITS) and the rDNA locus of a closely-related Vinca species (NCBI taxon ID 141628), over 92% of all reads were successfully classified (Figure 5). As expected, a large proportion of the reads generated with the 3.5 kB primer set is classified as host (*Vinca* sp.), while the new primer pairs did not generate such amplicons. In terms of species composition, the samples generated with the new primer pairs show similar abundances of the most abundant taxa with only slight variations. This demonstrates that our newly-designed primer pairs are indeed able to efficiently discriminate between the rDNA locus of fungi and the host plant and that they provide a similar reflection of the community composition.

For completeness, we also analyzed the data using the SILVA database, which includes a broader range of eukaryotes. For amplicons generated with primer pair 7 97% of the reads were classified as fungi, with primer pair 9 95%, and with primer pair 10 97%. Overall, none of the amplicons generated with the new primer pairs were classified as plant. Interestingly, a subset of amplicons was assigned to protist taxa, with the highest proportion, approximately 4.5%, observed in primer set 9. This indicates that while all primer pairs can successfully exclude the plant rDNA locus, primer pair 9 may be also good at targeting other microeukaryotes, which requires further investigation (Figure S3).

**Figure 5.**
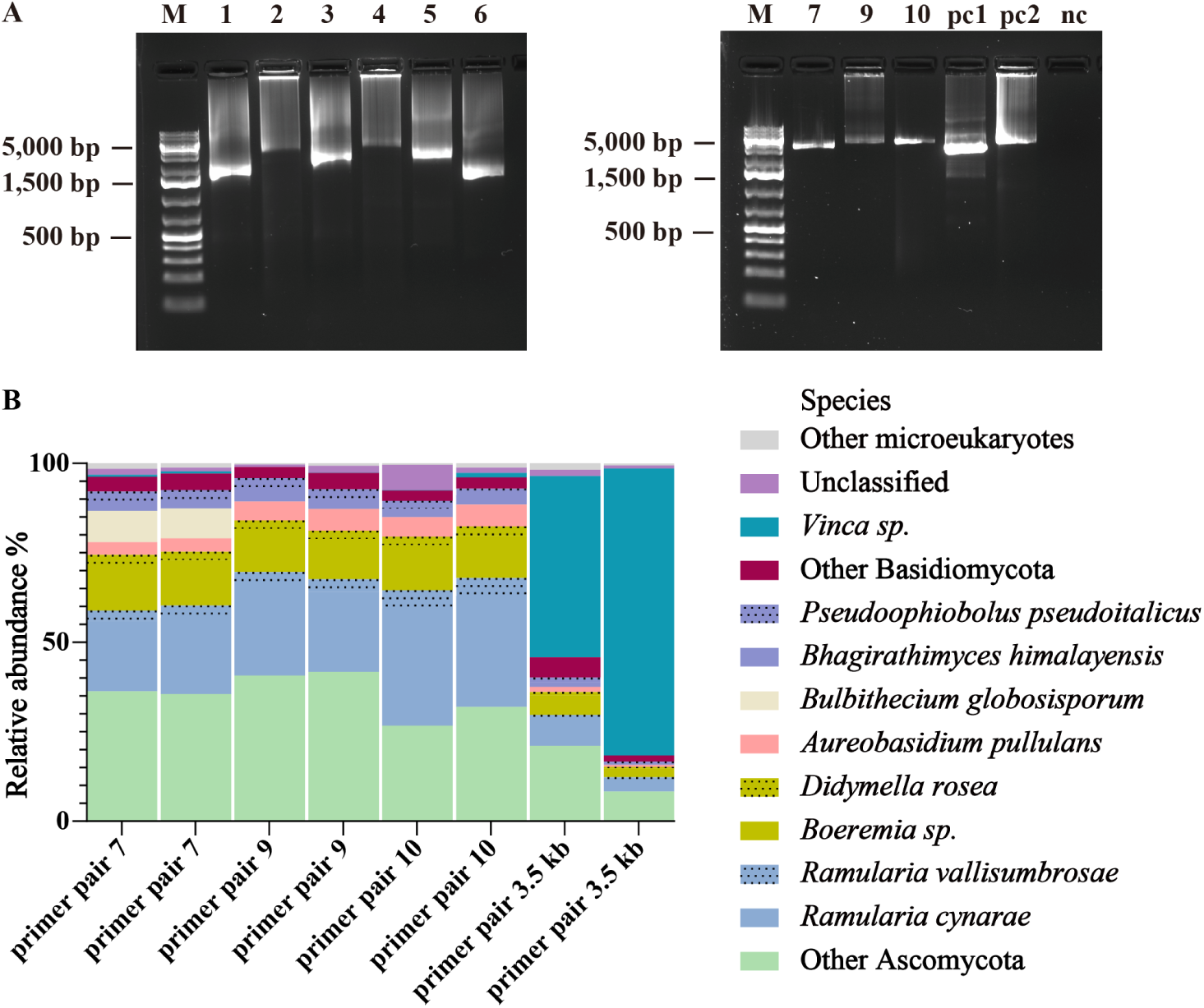
Metabarcoding of holobiont DNA of *V. minor*. (A) Gel electrophoresis of rDNA amplicons generated from gene specific primers including overhangs for PCR barcoding; Sample order in both panels: M: 1 kb plus DNA ladder; 1-10: amplicons generated with primer pairs 1-10 respectively; pc1: positive control generated with primer pair 3.5 kb; pc2: positive control generated with primer pair 10 with fungal mock community; nc: negative control (nuclease-free water). (B) Relative abundance of taxonomic groups classified based on the sequencing reads of amplicons generated from the genomic holobiont DNA of *V. minor* with the different primer pairs on species level (each primer pair has two replicates).

## Discussion

In molecular biology, the presence of non-target DNA (e.g., host DNA) can interfere with the detection and analysis of target DNA (e.g., microbial, pathogen, or environmental DNA). This is particularly problematic in fields like microbiome research, clinical diagnostics, and environmental DNA (eDNA) studies, where target DNA is often present in low abundance relative to other DNA. To address this issue, various blocking methods have been developed to selectively suppress the amplification or sequencing of non-target DNA. These techniques involve the use of chemically modified oligonucleotides, such as C3 spacers, locked nucleic acids (LNAs), or peptide nucleic acids (PNAs), which are designed to bind conserved regions of non-target DNA. The goal is to either direct the binding of blocking oligos to these modified sequences or to physically block polymerase elongation during PCR amplification. However, their effectiveness is often constrained by suboptimal binding affinity of the blocking oligos to non-target templates (Vestheim & Jarman 2008), the requirement for complex experimental optimization (Piñol *et al*. 2015), and limited cost-effectiveness (Azadnia *et al*. 2023). Given these limitations, the design of highly specific, discriminative primers could be a more practical and scalable alternative for minimizing non-target amplification.

In this study, we developed fungus-specific primers using a host-exclusive primer design workflow for the generation of long amplicons that almost entirely exclude plant DNA as a cost-effective alternative to the blocking methods. High-throughput sequencing of taxonomic marker genes (metabarcoding) is a widely used and powerful technique for biodiversity and microbial ecology. However, many taxa recovered from natural environments remain unidentifiable at the species level or even higher taxonomic levels when working with short reads (Tedersoo *et al*. 2020). In this case, long-read sequencing has the potential to overcome this problem. For instance, several of the six fungal isolates used for our mock community cannot be unambiguously identified solely based on ITS sequences (amplified with ITS1F and ITS4; data not shown) because they exhibit 100% sequence similarity with multiple taxa. For instance for the *Thyridium oculorum* isolate, ITS-based identification returned matches to both *Phialemonium* sp. and *Thyridium* sp., thus preventing a definitive taxonomic assignment. However, long-read sequencing results from this study showed that 92.15% ∼ 99.17% of all reads derived from this isolate were classified as *Thyridium oculorum*, which is consistent with the identification based on multi-locus classification. This highlights that long-read sequencing can provide superior classification results in metabarcoding studies.

There are, however, still common pitfalls that need to be tackled in the future. First, reducing bias in the relative abundance of sequence variants should be a primary objective in enhancing amplicon sequencing methods. It is well established that even PCR primers designed for broad coverage tend to amplify some phylogenetic subgroups more effectively than others. For instance in this study, the inferred abundance of *Trametes versicolor* in the mock community was consistently lower than the expected ratio, which might be attributed to amplification efficiencies of the used primer sets. In the experiments with holobiont DNA, we found that the overall community profiles were fairly consistent across primer pairs, though some minor differences were observed in the dominant taxa. This highlights the importance of evaluating primer performance in both controlled and real-world contexts when selecting appropriate primers for a given study.

Second, the accuracy of taxonomic classification is highly dependent on the reference database employed. Currently, these databases mainly rely on parts of the rDNA locus. The SILVA database includes near full-length sequences of the small and large subunits of the rRNA operon for both prokaryotes and eukaryotes. However, these sequences are insufficient for amplicons spanning the ITS region and generally allow identification only up to the genus level. Despite this limitation, the SILVA database enables the identification of non-fungal sequences, such as those from Chloroplastida and Metazoa (Quast *et al*. 2012). Conversely, for classification of fungi, the NCBI RefSeq Target Loci database seems to outperform the SILVA database. This superior performance is due to NCBI’s focus on the RefSeq database, which includes ribosomal DNA loci (16S, 18S, 28S rDNA genes, and ITS region) and allows for species-level identification, offering better accuracy and resolution in taxonomic classification of microbes. The limitation of this database, however, is that it is focused on archaea, bacteria and fungi, and does not include plants or other micro- and macro-eukaryotes. Therefore, reads originating from such eukaryotes will either not be classified at all or incorrectly classified as fungi. Furthermore, the fungal tree of life is still rather sparsely sampled for molecular barcodes deposited in the RefSeq database. Thus, significant efforts are still needed to improve the database to enhance its completeness and accuracy. This could involve developing databases tailored for eukaryotic full-length rDNA operons (Krabberød *et al*. 2025) or customizing databases to meet specific research needs.

## Conclusions

In the current study, we developed a host-exclusive primer design workflow, which is centered around a new computation tool, *mbc-prime*, to design discriminative metabarcoding primers with the option to exclude the DNA amplification of non-target taxa, for instance the host in host-associated microbiomes. We demonstrated the use of *mbc-prime* when targeting endophytic fungi associated with the medicinal plant *V. minor*. Initially, we evaluated our custom-designed primers *in silico*, demonstrating high coverage of fungal lineages with low coverage to Chloroplastida. Subsequent evaluation using a mock community revealed that the relative abundance of amplicons accurately reflected the community composition of the mock community. Moreover, when applied to plant holobiont samples, our primers successfully targeted a broad range of endophytic fungi while almost completely excluding the plant genome. This specificity has the potential to shed light on the roles of endophytic fungi in plant health and ecosystem dynamics. In addition, this demonstrates that our workflow, incorporating the *mbc-prime* tool, can effectively be used to study the composition of complex host-associated microbiomes. Furthermore, the primer design workflow developed in this study is universally applicable for designing discriminative primers targeting particular phyla, offering a user-friendly and practical solution for researchers in microbiome or other biological systems studies.

## Acknowledgments

We thank the Center for Information Technology of the University of Groningen for their support and for providing access to the Peregrine and Hábrók high-performance computing cluster. KH is grateful for financial support from the Federation of Biochemical Societies through the FEBS excellence award. THe and XL are supported by scholarships 202006550001 and 202106550001 from the China Scholarship Council. Figure 1 was created in BioRender (He, T. (2024) https://BioRender.com/d86p488).

