## Supplementary File 1 for "Host-Exclusive Primer Design Enables Effective Metabarcoding via Nanopore Long-Read Sequencing"

**Note S1**. Step-by-step tutorial for using mbc-prime.

**Note S2.** Step-by-step tutorial for building a Kraken2 custom database.

**Table S1**. Metadata for different primer combinations.

**Table S2** Metadata for different primer combinations as calculated by the OligoAnalyzer™ Tool.

**Table S3**. Fungal isolates used and their contribution to the mock community by pooling of rDNA at equimolar concentrations.

**Table S4**. Read statistics after de-multiplexing of sequenced amplicons generated from the mock community with various primer pairs.

**Table S5**. Read statistics after de-multiplexing of sequenced amplicons generated from the *V. minor* holobiont genomic DNA with different primer pairs, and classification results.

**Fig. S1.** The design of degenerate primers illustrated in the alignment of fungi, plants, and primer sequences.

**Fig. S2**. Metabarcoding of a fungal mock community classified with the NCBI RefSeq Targeted Loci database (ncbi_16s_18s_28s_ITS).

**Fig. S3.** Metabarcoding of holobiont DNA of *V. minor* classified with the SILVA database.

### Supplementary Notes

**Note S1.** Step-by-step tutorial for using *mbc-prime*
The following commands provide in a nutshell an example workflow for doing a primer design analysis with mbc-prime:

1) Move groups of sequences from database files into the same file (in the right order! inclusive group first, exclusive group second)
   $ seqkit grep -nrp Fungi databasefile1.fa >> sequences.fa
   $ seqkit grep -nrp Gentianales databasefile2.fa >> sequences.fa
2) To run mafft alignment; set as your input file sequences.fa, your output is e.g. alignment.aln, keep the input order
   $ bash ~/MAFFT/mafft-linux64/mafft.bat
3) To view alignment with Aliview (only if you want to check the alignment manually)
   $ aliview alignment.aln
4) To run mbc-prime (on an alignment file where the group you intend for primer design is 118 sequences big, and the desired minimum score is 0.8)
   $ ~/mbc-prime/mbc-prime -t 118 -s 0.8 alignment.aln
5) For analysis, it is most convenient to save this to csv (which can be opened in a spreadsheet editor):
   $ ~/mbc-prime/mbc-prime -t 118 -s 0.8 alignment.aln >> testalignment.csv
6) For more information, visit the mbc-prime repository on github, or view:
   $ ~/mbc-prime/mbc-prime -h

**Note S2.** Step-by-step tutorial for building a Kraken2 custom database.

```
# Step 1: Inspect FASTA headers
echo "Checking FASTA headers in f42_modified.fasta:"
grep "^>" f42_modified.fasta
# Step 2: Modify FASTA headers to include taxonomic ID (example: taxid 331006)
awk '/^>/ {print $1 "|kraken:taxid|331006"} !/^>/ {print}' f42_reference.fasta > f42_modified.fasta
# Step 3: Remove undesired taxid strings (if needed)
# Example: remove incorrect or outdated taxid entries
sed -i 's/|taxid|331006//' f42_modified.fasta
# Step 4: Download minimal Kraken2 taxonomy (skip mapping to NCBI taxids)
kraken2-build --download-taxonomy --db custom_db --skip-maps
# Step 5: Add custom reference genomes (FASTA) to the database library
kraken2-build --add-to-library f42_modified.fasta --db custom_db
# Step 6: Build the custom Kraken2 database
kraken2-build --build --db custom_db
# Step 7: Classify amplicon or metagenomic reads
kraken2 --db custom_db --threads 8 \
  --report output.report \
  --output output.kraken2 \
  --use-names input.fastq
# Step 8: Build Bracken database (for abundance estimation)
bracken-build -d custom_db -l 2000 -t 8
# Step 9: Run Bracken to estimate relative abundances at species level
bracken -d custom_db -i output.report -o bracken_report.txt -r 2000 -l S
# Step 10: Inspect the custom Kraken2 database structure
kraken2-inspect --db custom_db | less
# Optional: Clean up entire database (use with caution!)
# kraken2-build --clean --db custom_db
```

Supplementary Tables

**Table S1.** All primer pairs used in this study.

| Pair | Primers (fwd/rev) | Tm (°C) | Product size[a] |
|---|---|---|---|
| 1 | mbc-2479-F/LR5-Fung | 50 | 1.4-1.8 kb |
| 2 | mbc-2479-F/mbc-3159-R | 50 | 2.9-3.3 kb |
| 3 | mbc-2479-F/mbc-2148-R | 49 | 2.1-2.4 kb |
| 4 | mbc-1840-F/mbc-3159-R | 49 | 3.2-3.5 kb |
| 5 | mbc-1840-F/mbc-2148-R | 50 | 2.2-2.7 kb |
| 6 | ITS1catta/LR5-Fung | 55 | 1.3-1.6 kb |
| 7 | mbc-136-F/LR5-Fung | 50 | 3.1-3.4 kb |
| 8 | mbc-136-F/mbc-3159-R | 49 | 4.3-4.9 kb |
| 9 | mbc-136-F/mbc-2148-R | 50 | 3.6-4.0 kb |
| 10 | mbc--1220-R/mbc-3159-R | 46 | 3.7-5.0 kb |
| 3.5kb | SSU-Fngs-F/LR5-R | 55 | 3.5 kb |

Tm: The optimized annealing temperatures are based on the LongAmp hot start polymerase

a: The product size is estimated by the Primer-BLAST (https://www.ncbi.nlm.nih.gov/tools/primer-blast)

**Table S2.** Metadata for different primer combinations as calculated by the OligoAnalyzer™ Tool. Data layout for the table is illustrated below.

| Rv ↓ | Fw → | mbc-136-F | | mbc-1220-F[a] | | mbc-1840-F | | mbc-2479-F | |
|---|---|---|---|---|---|---|---|---|---|
| LR5-Fung | | 6 | -7.05 | 11 | -5.09 | 7 | -5.19 | 2-7 | -6.66 |
| | | 51 | 3100-3400 | 46 | 2300-2600 | 50 | 1600-2200 | 50°C | 1400-1800 |
| mbc-3159-R | | 2 | -4.13 | 3 | -5.57 | 1 | -7.13 | 1-6°C | -5.19 |
| | | 49 | 4300-4900 | 46 | 3700-5000 | 49 | 3200-3500 | 49°C | 2900-3300 |
| mbc-2148-R[b] | | 2-4 | -7.05 | 7-9 | -5.09 | 3-5 | -4.67 | 0-3°C | -7.26 |
| | | 51 | 3600-4000 | 46 | 2800-3300 b | 50 | 2200-2700 | 50°C | 2100-2400 |

a The original mbc-1220-F generated from mbc-prime was two bases (GA) shorter at the 5' end. We added these bases to the final primer to increase the melting temperature.

b In the final mbc-2148-R primer, we removed one base (C) at the 5' end compared to the original mbc-prime output in order to decease the melting temperature.

c Tm is calculated for LongAmp hot start 2x master mix.

d ΔG (Gibbs Free Energy): This value represents the thermodynamic stability of a structure. It is expressed in kilocalories per mole (kcal/mol) and determines whether a particular oligonucleotide structure (like a hairpin or dimer) is thermodynamically favorable. If ΔG less than -9 kcal/mol, this indicates a strong likelihood of primer dimer formation.

Data layout for each primer combination:

| | |
|---|---|
| Difference in Tm[c] (°C) | ΔG for heterodimerization[d] (kcal/mol) |
| Annealing temperature (°C) | Approx length of PCR product (bp) |

**Table S3.** Fungal isolates used and their contribution to the mock community by pooling of rDNA at equimolar concentrations.

| Phylum | Class | Order | Family | Genus | Species[a] | % of mock community | WGS-based identification[b] |
|---|---|---|---|---|---|---|---|
| **Basidio-mycota** | Agarico-mycetes | Polyporales | Polyporaceae | Trametes | ***Trametes versicolor*** | 16.66 | *Trametes versicolor*[c] |
| **Asco-mycota** | Sordario-mycetes | - | Thyridiaceae | Thyridium | ***Thyridium oculorum*** | 16.66 | *Thyridium* sp.CBS 149464[d] |
| **Asco-mycota** | Dothideo-mycetes | Pleosporales | Melanommataceae | Camposporium | ***Camposporium septatum*** | 16.66 | *Aposphaeria corallinolutea*[e] |
| **Asco-mycota** | Leotio-mycetes | Helotiales | - | Cadophora | ***Cadophora luteo-olivacea*** | 16.66 | *Cadophora luteo-olivacea*[f] |
| **Asco-mycota** | Dothideo-mycetes | Pleosporales | Cucurbitariaceae | Neocucurbitaria | ***Neocucurbitaria salicis-albae*** | 16.66 | *Neocucurbitaria salicis-albae*[g] |
| **Asco-mycota** | Sordario-mycetes | Hypocreales | Nectriaceae | Cosmospora | ***Cosmospora micropedis*** | 16.66 | *Cosmosporella olivacea*[h] |

a Classification results of amplicon sequencing in this study;

b Identification based on WGS information;

c Identification based on 4481 single-copy orthologous genes across 13 species using OrthoFinder;

d Identification based on 3532 single-copy orthologous genes across 7 species using OrthoFinder;

e Multilocus identification based on rbp2, tef1, LSU, SSU, ITS;

f Multilocus identification based on *rbp2*, *tef1*, *tub2*, LSU, SSU, ITS;

g Multilocus identification based on *rbp2*, *tef1*, LSU, SSU, ITS;

h Multilocus identification based on *tub2*, LSU, ITS.

**Table S4.** Read statistics after de-multiplexing of sequenced amplicons generated from the mock community with various primer pairs.

| Primer set | Number of reads after basecalling | Number of reads after trimming* | Mean length (bp) | Mean Qscore | Total yield (Mb) |
|---|---|---|---|---|---|
| 7 | 24,963 | 19,887 | 3,124 | 18.1 | 62.1 |
| | 35,782 | 29,996 | 3,125 | 18.2 | 93.7 |
| 8 | 19,008 | 11,776 | 4,498 | 18.0 | 53.0 |
| | 33,468 | 26,016 | 4,537 | 18.1 | 117.0 |
| 9 | 32,723 | 26,531 | 3,659 | 18.1 | 97.1 |
| | 39,945 | 33,761 | 3,644 | 18.3 | 123.0 |
| 10 | 30,433 | 22,166 | 3,812 | 18.1 | 84.5 |
| | 23,057 | 18,735 | 3,825 | 18.1 | 71.7 |
| 3.5 kb | 27,129 | 18,767 | 2,745 | 17.9 | 51.5 |
| | 20,945 | 16,899 | 2,743 | 17.9 | 46.4 |

* Reads were trimmed in the range of 2 kb – 6 kb.

**Table S5.** Read statistics after de-multiplexing of sequenced amplicons generated from the *V. minor* holobiont genomic DNA with different primer pairs, and classification results.

| Primer pair | Number of reads (after basecalling) | Number of reads (after trimming[a]) | Mean length (bp) | Mean Qscore | Total yield (Mb) | Classification proportion (%) |
|---|---|---|---|---|---|---|
| 7 | 61,766 | 49,377 | 3,056 | 18.4 | 151 | 99.90 |
| | 81,611 | 68,262 | 3,235 | 18.4 | 208 | 99.99 |
| 9 | 36,213 | 29,552 | 3,488 | 18.2 | 103 | 100.00 |
| | 72,599 | 55,396 | 3,499 | 18.2 | 194 | 99.99 |
| 10 | 74,663 | 64,301 | 3,682 | 18.2 | 237 | 99.94 |
| | 78,052 | 66,834 | 3,652 | 18.3 | 244 | 99.99 |
| 3.5 kb | 76,702 | 63,370 | 2,745 | 18.2 | 174 | 99.99 |
| | 84,937 | 75,240 | 2,772 | 18.2 | 209 | 99.99 |
| PC[b] | 31,522 | 3,648 | 3,660 | 18.2 | 13.4 | 99.97 |
| | 22,274 | 2.979 | 3,653 | 18.2 | 10.9 | 100.00 |
| NC[c] | 1,248 | 21 | 2,734 | 16.3 | 5.74e-08 | - |

a Reads were trimmed in the range of 2kb – 6kb;

b Positive control with mock community as input along the whole experimental procedure;

c Negative control with only nuclease-free water as input along the whole experimental procedure.

Supplementary Figures

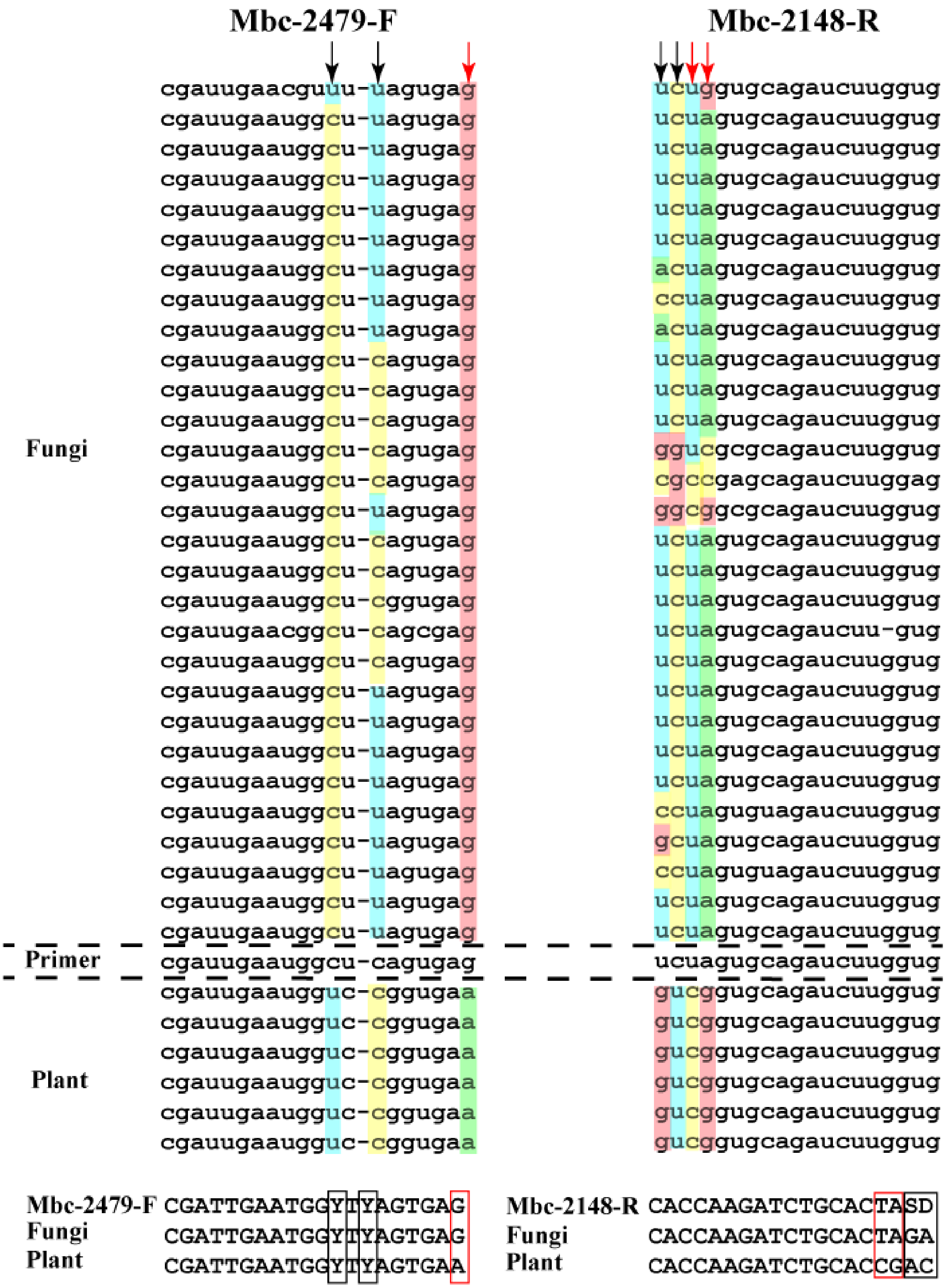


**Figure S1.** The design of degenerate primers illustrated in the alignment of fungi, plants, and primer sequences. Black arrows indicate degenerate positions and red arrows indicate discriminative sites on primer mbc-2479-F and mbc-2148-R. Y = C/T, S = C/G, D = A/G/T.

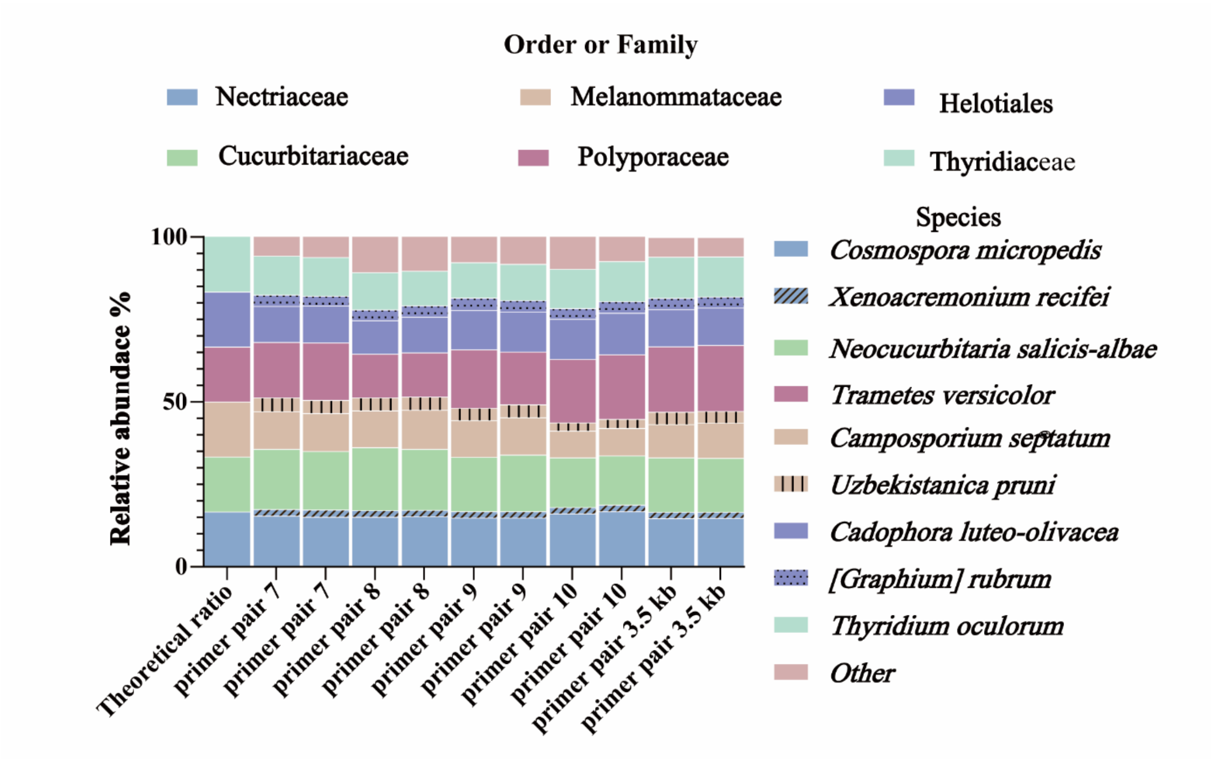


**Figure S2.** Metabarcoding of a fungal mock community classified with the NCBI RefSeq Targeted Loci database (ncbi_16s_18s_28s_ITS). Relative abundance of taxonomic groups classified based on the sequencing reads of amplicons generated from the mock community with the different primer pairs (each primer pair has two replicates). Species belonging to the same family are displayed in the same color with different fill patterns. All primer pairs reproduce the overall structure of the mock community, albeit with different biases for different groups.

Among the six isolates, four were correctly identified at species level (Figure S2, Table S3). The isolates of *Aposphaeria corallinolutea* and *Cosmosporella olivacea*, which lack species representatives in the RefSeq database, were primarily assigned to two close relatives in the database, *Camposporium septatum* and *Cosmospora micropedis*, respectively. *Cosmosporella* and *Cosmospora* are genera in the family *Nectriaceae* (Huang *et al.*, 2018), while *Aposphaeria* and *Camposporium* are members of the Pleosporales (Hyde *et al.*, 2020). A minor fraction of the reads was moreover classified as *Uzbekistanica pruni*, which also belongs to the family of *Melanommataceae* and order of Pleosporales.

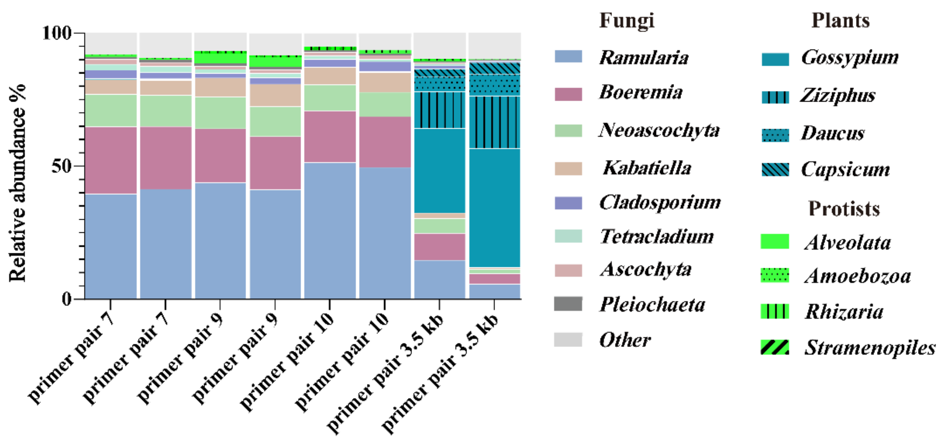


**Figure S3.** Metabarcoding of holobiont DNA of *V. minor* classified with the SILVA database. Relative abundance of taxonomic groups classified based on the sequencing reads of amplicons generated from the genomic holobiont DNA of *V. minor* with the different primer pairs on species level (each primer pair has two replicates).
